# Multiplexed Single Ion Mass Spectrometry Improves Measurement of Proteoforms and Their Complexes

**DOI:** 10.1101/715425

**Authors:** Jared O. Kafader, Rafael D. Melani, Kenneth R. Durbin, Bon Ikwuagwu, Bryan P. Early, Ryan T. Fellers, Steven C. Beu, Vlad Zabrouskov, Alexander A. Makarov, Joshua T. Maze, Deven L. Shinholt, Ping F. Yip, Danielle Tullman-Ercek, Michael W. Senko, Philip D. Compton, Neil L. Kelleher

## Abstract

A new Orbitrap-based single ion analysis procedure is shown to be possible by determining the direct charge on numerous measurements of individual protein ions to generate true mass spectra. The deployment of an Orbitrap system for charge detection enables the characterization of highly complicated mixtures of proteoforms and their complexes in both denatured and native modes of operation, revealing information not obtainable by traditional measurement of an ensemble of ions.

---

For decades, mass spectrometry has used ions to measure the mass-to-charge ratio of molecules once lifted into the gas phase^1^. Denatured and native electrospray ionization of intact proteins and their complexes pose many complications due to sample heterogeneity and large charge state envelopes in the *m/z* domain. To simplify analysis, Charge Detection Mass Spectrometry (CDMS) ^2-6^ has enabled the generation of true *mass* spectra with the direct readout of an ion’s integer charge value (e.g. 35+, 34+, etc.). Here, we bring the commercially available Orbitrap mass analyzer into the CDMS universe to multiplex and regularize this approach for creation of mass spectra on extremely complex samples. The approach is shown to be general for measuring complex proteoform mixtures and their complexes, covering a range of masses from 8 kDa to 3.2 MDa, with Orbitrap-based harmonic Charge Detection Mass Spectrometry (hCDMS) capable of precise mass determination where ensemble measurements fail or require prior separation.

Previously, single ion sensitivity has been demonstrated using an Orbitrap mass analyzer^7, 8^, analogous to linear geometry trapping instruments more frequently used for CDMS. In 2018, we used an Orbitrap analyzer to acquire one-at-a-time measurements on single ions, and showed that >20-fold increases of resolving power over ensemble measurements were possible by centroiding *m/z* values of individual ions^9^. Here, we extend this methodology substantially with a complete workflow for CDMS that we call hCDMS (**Fig. 1**) to directly measure the charge of individual ions. Further, we show this method to be fully established and functional for the simultaneous measurement of 200 individual ions.

**Figure 1.**
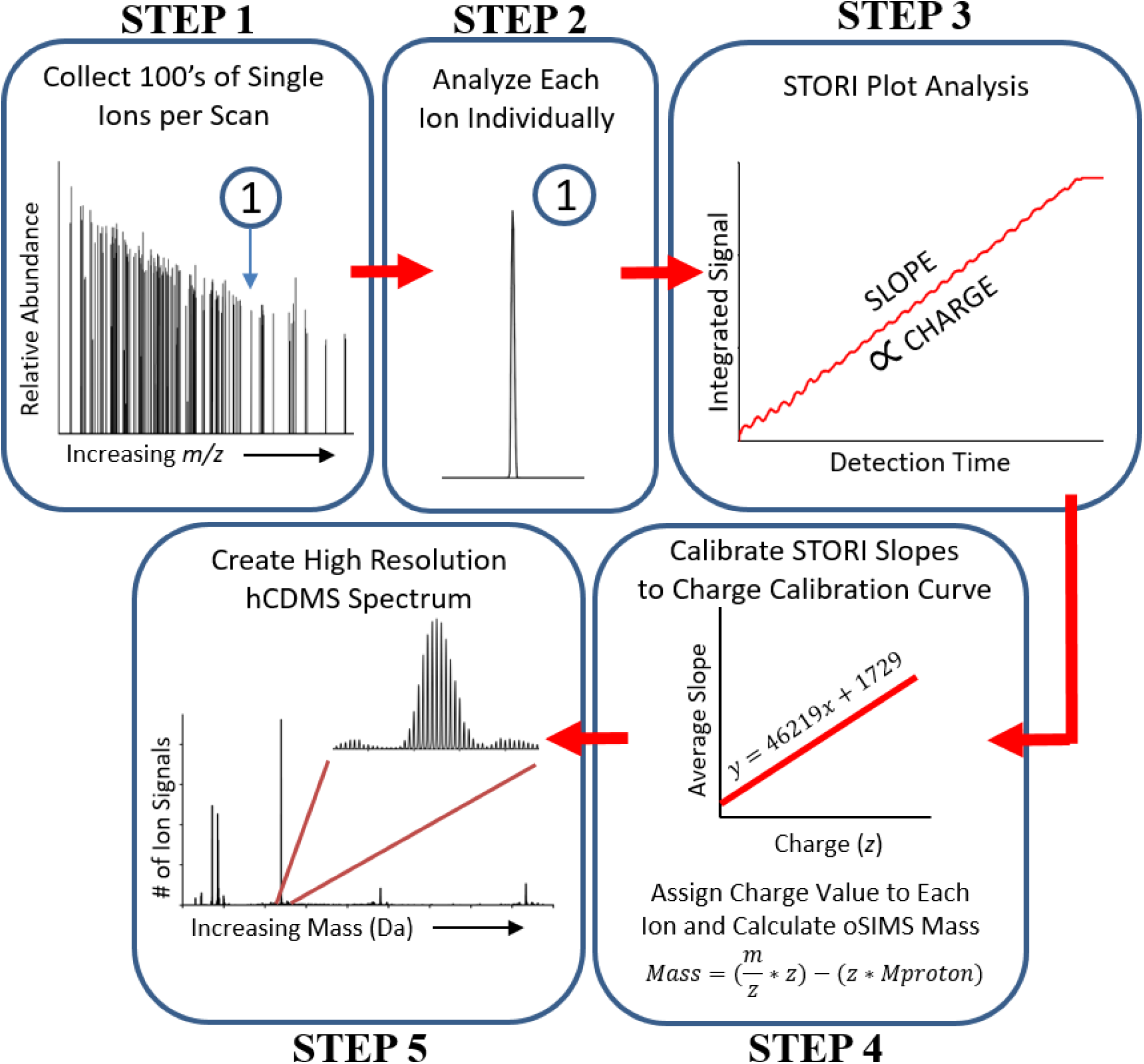
The five step workflow for the hCDMS process, which includes data acquisition and post-acquisition processing steps that culminate in the plotting of an hCDMS spectrum (Step 5). A detailed explanation of each of these five steps is found in the accompanying Online Methods section.

In Step 1 **(Fig. 1**) of the hCDMS approach, ∼120 ions are observed per acquisition in a random-style trapping event. Parallelizing ion observation to >100 ions per acquisition decreases the data acquisition burden by ∼100-fold over current CDMS techniques^10^. Due to the low ion count needed for hCDMS, protein solutions can be diluted down by ∼three orders of magnitude (into the high pM to low nM range). In Step 2 (**Fig. 1**), the frequency of each ion signal is determined and analyzed independently. At this processing stage, precise information for each ion including frequency, intensity, and *m/z* values is established. Step 3 (**Fig. 1**) determines the ion’s signal strength using a data plotting and analysis process that assesses the current induced by an ion on the Orbitrap detection electrodes as a function of acquisition time. For simplicity we call this signal strength determination the “STORI” process^11^ with the slope of a STORI plot being proportional to the charge of the ion (Step 3, **Fig. 1**). In Step 4, the charge of the ion is determined by a slope-to-charge calibration function. The STORI slope of an ion with unknown charge is assigned the closest integer charge state on the calibration function, determined just once for each instrument (two different Q Exactive-style instruments were used in this study). Using the integer charge (*z*) and mass-to-charge ratio (*m/z*), it is possible to determine the mass of each ion and produce a spectrum in the true *mass* domain with quite different spectral properties and increase resolution via centroiding and binning single ion signals^9^. As a result of our charge assignment algorithm, further explained in the Online Methods, charge misassignments do occur at the 1-4% level of the total ion count in hCDMS spectra; these artifacts are recognized readily as satellite peaks due to the ±1 error in charge state assignment. Further experimental optimization and software development is currently underway to completely eliminate charge misassignment. The hCDMS process further uses validated raw data reduction and a reduced DC offset for the Orbitrap center electrode (−5 to −1 kilovolt) enabling greater than 70% of the ions to survive the ∼2-4 second detection period (**Supplemental Fig. 1**). A more in-depth description of these steps is available in the Online Methods. We now focus on the application of the method to complex proteoform mixtures and detection of large complexes via native hCDMS.

Initial experiments on a mixture of intact protein standards from 8-47 kDa highlighted an immediately apparent set of differences in hCDMS readout from conventional ensemble mass spectrometry analysis. Basic figures of merit for these standards are shown in **Supplemental Fig. 2c**, including resolving power over one hundred thousand (even at high mass) and usual low ppm errors (< 5 ppm errors in **Supplemental Table 1**). Note that hCDMS produces simple spectra with an x-axis of mass, removing the normal requirement for the inference step to convert *m/z* to mass. The remarkably clean baselines of hCDMS spectra proved highly insensitive to partially desolvated ions (**Supplemental Fig. 2a vs. 2b**), which extended the dynamic range for intact mass determination even when complexity was high (*vide infra*). The strict filtering criteria during STORI slope determination removed non-stable ion signals stemming from desolvation, ion fragmentation in the Orbitrap analyzer (such as neutral loss), and intermittent signals produced from electronics (electronic noise). In essence, hCDMS transforms the instrument into an actual MASS spectrometer, removing the normal requirement for the inference step in conventional mass spectrometry.

We then used hCDMS to analyze a mixture of heavy and light chains resulting from disulfide reduction of an IgG_1_ (**Supplemental Fig. 3**). Here, the removal of non-stable ion signals yielded highly sensitive hCDMS spectra of species and adduct identification that was not possible in ensemble measurements. Ultimately, these mass domain readouts greatly simplify data analysis for non-experts.

To stress-test the hCDMS process, it was compared to standard direct infusion with automatic gain control (AGC) set to 1×10^6^ charges/spectrum for high complexity samples like the entire <30 kDa range of the human proteome from HEK293 T-Cells fractionated by GELFrEE technology^12^. A silver-stained SDS-PAGE gel (inset of **Fig. 2a**) shows the range of proteins that were analyzed by both standard MS and hCDMS approaches (**Fig. 2a** vs. **2b**). Although no protein masses could be determined using standard MS (**Fig. 2a**), hCDMS (**Fig. 2b**) detected over 500 proteoform masses that were assigned using the intact mass tag approach^13, 14^ and referencing prior top-down proteomics data where tandem MS (LC-MS/MS) was used to create a list of high quality proteoforms present in the Human Proteoform Atlas^15^. A comparable number of ions (∼10 million) was utilized to generate the ensemble and hCDMS spectra. While all identified proteoforms via hCDMS analysis are listed in **Supplemental Table 1**, small insets in **Fig. 2b** illustrate 17 identified proteoforms that are listed in **Fig. 2c**. In addition, isotopic distributions for higher mass proteoforms between 20 – 25 kDa illustrated in the red box on **Fig. 2b** have not previously been identified and characterized by prior LC-MS/MS analysis. For conventional mass spectrometry, large proteins (>20 kDa) are burdened with charge state distributions consisting of dozens of visible states that a fixed number of ions are appropriated between. hCDMS, with the high intra-spectral dynamic range of ∼2,000 clearly identifies these high mass features usually not detectable in lower dynamic range traditional *m/z* analysis. hCDMS reduces charge state ambiguity and consolidates each charge state distribution into one mass channel, increasing detection capabilities for larger mass species in mixtures with extreme complexity.

**Figure 2.**
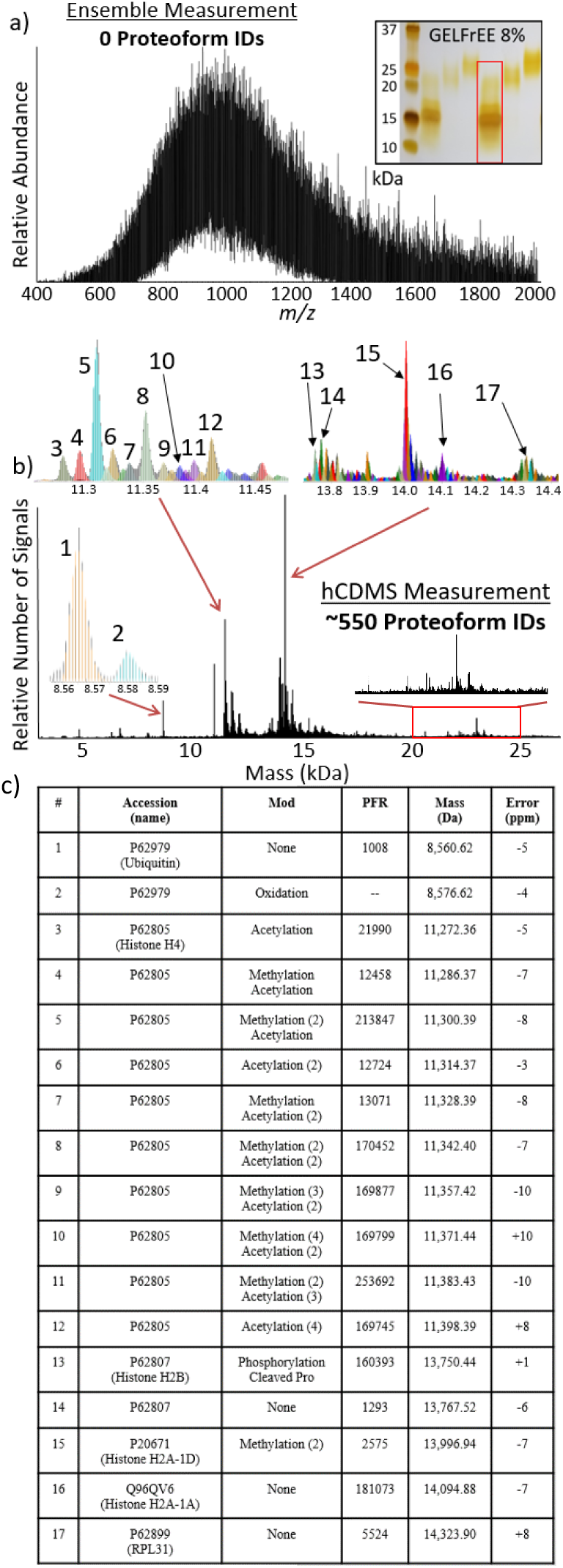
Comparison of number of proteoform assignments possible using either conventional spectral acquisition with and ensemble of ions in a population (a) or the hCDMS process (b). This comparative analysis involved directly infusing a mixture of 0 - 30 kDa proteins created by fractionation of whole extracts from HEK-293 human cells. The inset in panel (a) shows an analytical, silver-stained gel, with the fraction analyzed highlighted with a red rectangle. The insets in panel (b) showcase a subset of modified proteoforms from the over 500 identified proteoforms listed in (c) along with accession numbers, post-translational modifications, Proteoform Record numbers (PFR), monoisotopic masses, and experimental error (in ppm). The 20-25 kDa region highlighted with a red rectangle in panel (b) corresponds to proteoforms of higher mass proteins in the complex mixture previously unidentified using LC-MS/MS for top-down proteomics.

Another major advantage of hCDMS is the ability to work with samples containing large protein complexes that are lifted into the gas phase using native electrospray ionization (so called native MS)^16^. Pressing hCDMS further, we examined standard protein complexes^17-19^ and also applied it to engineered virus-like particles (VLP). As demonstrated by **Supplemental Fig. 4**, protein complexes including pyruvate kinase (**Supplemental Fig. 4a, b**, and **c**) and GroEL (**Supplemental Fig. 4d, e, and f**) with well resolved charge states validate hCDMS implementation for native complexes. The charge state distributions from hCDMS match those determined using standard MS, and the relative intensity of the charge assignments mirror their corresponding spectra in the *m/z* domain. Accurately assigned hCDMS single ion charge states produce very defined mass peaks for pyruvate kinase (231.9 kDa) and GroEL (801.2 kDa) with high resolution at these large mass values (650 – 800). Further, minor species of the pyruvate kinase complex formed from loss of a C-terminal lysine in one or two of the subunits within the tetramer (**Supplemental Fig. 4b**) are easily recognized.

The single-ion analysis advanced here was conducted on two virus-like particles (VLPs) engineered from viral capsids carrying varying amounts of DNA and mRNA cargo loads after heterologous production in *E. coli*. Wild-type (WT) or mutant (MINI) versions of these VLPs were constructed from the MS2 wild-type coat protein or a Ser37Pro mutant, respectively. Recently, these particles have been investigated through electron microscopy and X-ray crystallography to determine capsid structure^20-24^. Briefly, WT and MINI VLPs are composed of exactly 180 and 60 copies of the 13.7 kDa coat protein (CP), generating theoretical masses of 2.47 and 0.82 MDa, and particles with 27 and 17 nm diameters. In turn, cavity volumes can be calculated around 10,300 and 2,600 nm^3^, assuming an approximate spherical nature of the VLPs. The WT and MINI monomer amino acid composition was verified through MS analysis in denatured mode **(Supplemental Fig. 5)**. Although the structure of the VLPs is established, characterization of cargo such as therapeutic proteins and RNA would be highly valuable to quickly determine the stoichiometry and loading of drug-delivery within the particles.

To probe composition and cargo of the VLPs, hCDMS data for the WT and MINI particles were compared to each other and contrasted with data from standard MS data produced in the *m/z* domain (**Fig. 3c,d** and **Fig. 3e,f**). No visible charge states were present in the *m/z* domain spectra of either the WT (**Fig. 3c**) or MINI (**Fig. 3d**) species. The lack of distinct charge states in the native MS data was attributed to the molecular heterogeneity of varying lengths of DNA and mRNA^22^. Given this molecular heterogeneity and no distinct charge state peaks present in the *m/z*-domain spectra, no information regarding mass of the VLP plus cargo could be discerned. This is an often-encountered situation in obtaining size distributions of highly complex samples in mass spectrometry and the reason other methods like gel electrophoresis and light scattering are used to provide broad molecular weight estimations.

**Figure 3.**
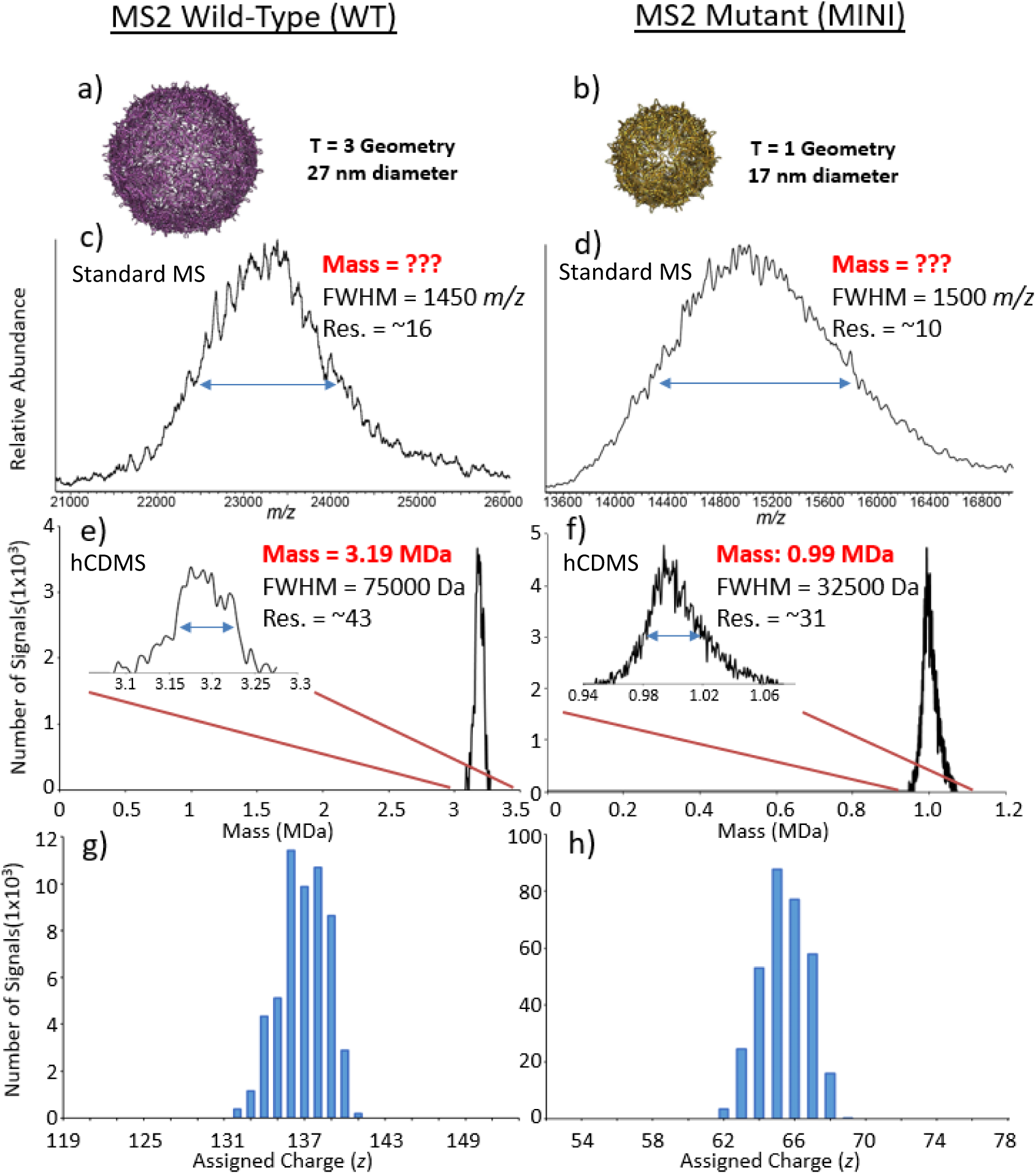
Crystal structure renderings (a,b) and mass spectra of assembled WT (c,e,g) vs. MINI (d,f,h) MS2 virus-like particles, with approximate masses and capsid diameters of 3.2 vs. 1 MDa and 27 vs.17 nm, respectively. In panels (c) and (d), the standard MS spectra plotted in *m/z* space results in a loss of charge state resolution due to heterogeneity of cargo inside the capsid of the virus-like particles. However, the corresponding hCDMS spectra in panels (e) and (f) allow mass assignment to the distribution of particles even without resolution of the individual charge states (a normal requirement of conventional data produced by electrospray MS). The individual ions assigned to their underlying charge distributions are shown in panels (g) and (h). Mass and resolution (Res.) at full-width of the half-maximum (FWHM) of the peak heights are labeled beside each peak.

In contrast to standard native MS, the hCDMS spectra produced mass distributions of the WT (**Fig. 3e**) and MINI (**Fig. 3f**) species with masses of 3190 ±38 kDa and 990 ±16 kDa (mean ± s.d.). Assigned charge states for the set of single ions were centered around +137 and +67 for WT (**Fig. 3g**) and MINI species (**Fig. 3h**) with the expected Gaussian-like distributions. Determined hCDMS masses minus proposed capsid masses reveal WT and MINI cargo masses of approximately 0.72 ± 0.038 MDa and 0.17 ± 0.016 MDa, respectively. Although the length of the DNA base pairs and mRNA nucleotides that form the VLP cargo loads vary across a wide distribution of lengths, MINI VLPs having a much smaller volume incorporate shorter DNA and mRNA strands than their WT counterparts. DNA and mRNA lengths averaged 900 (±400) vs. 375 (±150) base pairs (bp) and 500 (±250) vs. 100 (±50) nucleotides (nt) for WT vs. MINI species^22^. The mass distribution for the WT particles determined via hCDMS (**Fig. 3e**) is consistent with the encapsulation of 1 average length DNA (900 bp) and mRNA (375 nt) strand together in the VLP (∼0.74 MDa). The WT mass distribution with a FWHM of only 75 kDa means that as a larger DNA strand is added as cargo, it is partnered with a smaller length mRNA component during production and self-assembly within the cytosol of *E. coli.* The mass distribution for the MINI particles determined via hCDMS (**Fig. 3f**) is consistent with VLP encapsulation of either 1 DNA strand or multiple smaller mRNA strands as cargo. The incorporation of the average length DNA strand (375 bp) matches the higher mass tailing feature around 1.06 MDa. In turn, the smaller VLP volume can contain multiple (∼5) of the smaller ∼100 nt mRNA strands, accounting for the majority of the MINI mass distribution. The ratio, 4.2, of the cargo masses 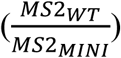 corresponds to the ratio of cavity volumes, (4.0) further supporting assignment of cargo content based on hCDMS data for these spherical WT and MINI VLPs. The measurement of complex masses through hCDMS, unobtainable via traditional mass spectrometry, reveals pertinent information to their composition and cargo.1

With increased dynamic range, resolution, clean baselines and intuitive data interpretation, hCDMS shows increased value for determination of mass values particularly as mixture complexity increases. Being an Orbitrap-based method, hCDMS provides a readily deployable technique for single molecular MS of denatured proteins and native complexes with masses ranging from 5 kDa to 3.5 MDa. The ability to accurately determine mass values directly from dilute and complex samples regardless of charge state overlap will greatly inform and expand the utility of mass spectrometric analysis of proteins and their assemblies in improved characterization of molecular complexity for both endogenous and synthetic biology.

## Online Methods

### Cell growth, lysis, and protein fractionation

Human embryonic kidney cells HEK-293 (ATCC CRL-1573) were grown on 15 cm plates in DMEM media (Life Technologies) supplemented with 10% fetal bovine serum (Life Technologies), containing 1% Penicillin/Streptavidin (Life Technologies), and they were incubated at 37°C temperature under 5% CO2 atmosphere. Triplicate of confluent plates containing 1 × 10^7^ cells were treated with 100 nM rapamycin or a DMSO control for 8 hours and harvested with 0.05% trypsin (Life Technologies) for 5 min. For CDMS experiments cells were not treated. Cells were pelleted at 300 x g for 10 min. and washed twice with PBS (Life Technologies). Cells were incubated with lysis buffer (1% w/v N-lauroylsarcosine, 20 mM Tris Base pH 7.5, 100 mM NaCl, and HALT Inhibitor Cocktail (Thermo Fisher Scientific) for 10 min. on ice, 750 units of Benzonaze nuclease (Sigma) was added, the solution was incubated for 15 min. at 37°C, and the protein content was estimated using Pierce BCA assay (Thermo Fisher Scientific). Aliquots of 400 µg of protein were precipitated for 30 min. at −80°C with 4 volumes of cold acetone, resuspended in GELFrEE Tris-Acetate Sample Buffer (Expedeon), and proteins were fractionated on a GELFrEE 8100 (Expedeon) system using an 8% cartridge according to manufacture procedures. Only the first fraction containing proteins up to 30 kDa was collected.

### Denatured mode top-down proteomics using LC-MS/MS

Fractions were methanol/chloroform/water precipitated^12^, resuspended in buffer A (5% acetonitrile, 94.8% water, and 0.2% formic acid), and submitted to LC-MS/MS. Samples were fractionated on a Dionex UltiMate 3000 LC system (Thermo Fisher Scientific) using an in-house packed PLRP-S (5 µm particle, 1000 Å pore size) (Agilent) column (75 µm ID × 25 cm length) after loading on a PLRP-S trap column (150 µm ID × 2 cm). The loading pump was operated at 3 µL/min., and the nano pump gradient flow rate was set at 300 nL/min. Buffer A and buffer B (5% water, 94.8% acetonitrile, and 0.2% formic acid) were used for LC separation under the following gradient: 5% B at 0 min., 15% B at 5 min., 55% B at 55 min., 95% B from 58 to 61 min., 5% B from 64 to 80 min. Column was kept at constant temperature (55°C), the outlet was online with a 15 µm i.d. electrospray emitter (New Objective) packed with 3 mm of PLRP-S resin, and attached to a nanoelectrospray ionization source built in-house. LC was online with an Orbitrap Fusion Lumos (Thermo Fisher Scientific) mass spectrometer operating in “protein mode” with 2 mTorr of N_2_ pressure. Transfer capillary temperature was set at 320 °C, ion funnel RF was set at 30%, and a 15V of source CID was applied. MS^1^ spectra were acquired at 120,000 resolving power (at 200 m/z), AGC target value of 500,000 charges/acquisition, 200 msec. of maximum injection time, and 4 µscans. Data-dependent top two MS^2^ method using 23 NCE for higher-energy collisional dissociation (HCD) was used to generate MS^2^ spectra that were acquired at 60,000 resolving power @ 200 *m/z*, with target AGC values of 500,000 charges/acquisition, 800 msec. of maximum injection time, and 4 µscans. Precursors were quadrupole isolated used using a 3 *m/z* isolation window, dynamic exclusion of 60 sec. duration, and threshold of 2×10^4^ intensity.

### Protein Identification

Raw files from LC-MS/MS experiments were searched against a UniProt *Homo sapiens* (Taxon ID: 9606, Proteome ID: UP000005640) protein database using TDPortal (http://galaxy.kelleher.northwestern.edu/), as previously described ^25, 26^. Proteoform search space was generated *in silico* allowing up to 11 PTMs or sequence variations per candidate proteoform. Two different searches types were performed in parallel: Absolute Mass search with precursor tolerance of 2.2 Da and fragments tolerance of 10 ppm, and a Biomarker search with 10 ppm precursor and fragments tolerance. Estimation of false discovery rate (FDR) at the protein entry and proteoform level was performed^27^. Proteoforms above a 10% FDR threshold were reported and used for GELFrEE fraction hCDMS spectrum annotation.

Proteoforms passing the FDR cutoff were transformed into theoretical +1 (M+H) isotopic distributions^27^. The hCDMS mass spectrum was also converted into singly-charged data for comparison. With the identified proteoforms serving as theoretical distributions, the hCDMS distributions were matched against the theoretical using an isotope fitting routine with a 25 ppm tolerance^28, 29^. Matches were loaded into TDValidator for visualization and manual validation (Proteinaceous, Inc.).

### Virus-like Particle Expression and Purification

Glycerol stocks of *E. coli* DH10B cells containing plasmids encoding for either the MS2 wild-type coat protein or the mutant (mini) variant were grown in 10 mL 2xYT overnight at 37°C. These cells were then subcultured into 1 L 2xYT to a starting OD600 of 0.05 and grown with shaking at 37°C. When the OD600 reached 0.5, protein expression was induced with 0.1% arabinose. Expression continued overnight for ∼ 16 h. After harvesting the cells by centrifugation, the cellular protein was extracted and VLPs were purified according to standard procedures using ammonium sulfate precipitation and fast protein liquid chromatography (FPLC)^24^.

### Additional Sample Preparation and MS Analysis

To produce the synthetic 4 protein sample mixture (**Supplemental Fig. 2**) ubiquitin, myoglobin, carbonic anhydrase, and enolase purchased from Sigma Aldrich were added together to produce a 2 ml solution with final protein concentrations of 20, 40, 50, and 90 nM, respectively. To produce the reduced version of the NIST antibody standard (**Supplemental Fig. 3**) 30 µL (300 µg) of stock solution (NIST) was mixed with 300 µL of 6 M GdHCl. 30 mM TCEP-HCl was added, the solution was incubated at 37°C for 1.5 h, and finally diluted to a 500 nM concentration. HEK293 GELFrEE fraction 1 (Fig. 1) was methanol/chloroform/water precipitated^12^, resuspended in HPLC grade water and run through a HiPPR (Thermo Fisher Scientific) detergent removal column, following the manufacture’s protocol. The fraction protein concentration was determined with a BSA protocol (Thermo Fisher Scientfic) and diluted to a 5 nM concentration. Before dilution all samples were desalted six times at 10,000 x *g* for 5 min. in 3 kDa Amicon Ultra centrifugal filters (Merck Milipore) with 100 mM ammonium acetate. Once desalted, all samples were diluted and electrosprayed under denaturing conditions in a 40% acetonitrile and 0.2% acetic acid solution.

Pyruvate kinase and GroEL protein complexes (**Supplemental Fig. 4**) were purchased from Sigma Aldrich and diluted to final concentrations of 1 µM. GroEL was precipitated as previously described^30^. Both protein complexes were desalted six times at 10,000 x *g* for 4 min. in 100 kDa Amicon Ultra centrifugal filters (Merck Milipore) wtih 100 mM ammonium acetate. Once desalted, both samples were diluted and electrosprayed under native conditions in 100 mM ammonium acetate.

Denatured samples with molecular weights less than 100 kDa were introduced into a Q Exactive Plus (Thermo Fisher Scientific) mass spectrometer with a custom nano electrospray source as described previously^31^ using +0.8 to 1.6 kV spray voltage. The mass spectrometer was modified to allow for a reduced Orbitrap central electrode voltage up to −1 kV. Native samples with molecular weights greater than 100 kDa were introduced into a Q Exactive Ultra High Mass Range (UHMR) (Thermo Fisher Scientific) mass spectrometer with a Nanospray Flex Ion Source (Thermo Fisher Scientific) with spray voltages between 1.4 to 2 kV. For both instruments acquisition rates were reduced down to 0.5 spectra/sec, corresponding to a 2 second detection period per acquisition event and the HCD-pressure was set between 0 to 0.5 (UHV Pressure < 5×10^-11^ torr) to reduce ion and background-gas collisions from a more typical setting of 1 (UHV Pressure ∼9×10^−11^ torr). Enhanced Fourier-Transform (eFT) was turned off, as it was not necessary to process single ion signals in this manner. A source temperature of 320°C, an in-source collision-induced dissociation (SID) value of 0 to 15 V, and an in-source trapping (UHMR) value of −50 to −150 V was optimized on a per species basis.

### Ion Collection and Data Acquisition of the hCDMS Process

The overall scheme describing hCDMS is shown in **Fig. 1**. Below is an extended technical description of this new process.

### Ion Collection (Step 1)

Similar to traditional CDMS techniques, hCDMS evaluates single ion signals to produce a true mass spectrum. However, Orbitrap CDMS analysis differs from traditional linear ion trap CDMS in the efficiency of single ion collection^2-6^. With the implementation of image current detectors our single ion definition is not confined to one ion per acquisition event, but one ion at a defined frequency per acquisition, meaning that multiple ions are analyzed per acquisition as long as they do not correspond to multiple ions at the same *m/z* value. To lower the number of ions entering the Orbitrap analyzer, denatured samples were diluted to concentrations between 5-50 nM, while samples of large complexes were analyzed by native electrospray in the 1-3 μM regime. In addition, Automatic Gain Control (AGC) was disabled and ion injection times were set to between 0.03 – 1 ms. In some cases the ion optics including the C-trap entrance lens were detuned to further reduce the number of ions entering the Orbitrap analyzer.

### Individual Ion STORI Slope and Centroid *m/z* Determination (Fig. 1, Steps 2 and 3)

Within each acquisition event, every detected ion was analyzed individually. Both the mass-to-charge ratio (*m/z*) and charge (*z*) were necessary to determine the mass of the ion in question. In FT-MS Orbitrap analysis, *m/z* was determined from the frequency of ion rotation around the central electrode. To determine the centroid *m/z* value, the apex of the profile peak was determined by a quadratic fit to the three most intense *m/z* points. The *z* was measured from the rate of the induced charge on the Orbitrap outer electrode, otherwise known as Selective Temporal Overview of Resonant Ions (STORI) as described in the accompanying STORI manuscript (included here as **Appendix 1**). Briefly, STORI analysis was the integration of the ion induced charge over the course of the time domain acquisition period at the specified single ion frequency value. The slope of the STORI plot increased at a constant rate until either the end of the transient acquisition event or ion disintegration within the Orbitrap analyzer. The STORI plot slope of a particular ion was proportional to the charge of the observed ion. **Supplementary Fig. 6** demonstrates the dependence depicting increasing STORI slopes for the +13 charge state of ubiquitin, the +25 charge state of myoglobin, and the +39 charge state of carbonic anhydrase. STORI slope is only dependent on the charge of the ion and not the *m/z* or mass of the species in question.

### Charge Determination and Slope Averaging Technique (Fig. 1, Step 4)

In order to accurately assign the unknown *z* of an ion with a known STORI slope value, each ion slope was calibrated to a charge calibration function. **Supplemental Fig. 7** illustrates 2 linear calibration functions created from isolated ions with known charge states analyzed with a Q Exactive Plus (red points/trace) and Q Exactive UHMR (blue points/trace). To produce the trace for the Q Exactive Plus various known charge states from denatured ubiquitin, myoglobin, carbonic anhydrase, and enolase were analyzed. To produce the trace for the Q Exactive UHMR various known charge states from natively electrosprayed carbonic anhydrase, monoclonal NIST antibody standard, pyruvate kinase, and GroEL were analyzed. Thousands of single ions from each known charge state were analyzed and their median STORI slope value was calculated (represented by each black point). Each calibration file was manually curated to ensure the STORI slopes utilized to calculate the average value did not include slopes from two ion signals at the same *m/z* value or slopes from contaminating species that overlap in *m/z* space. The points on the calibration were fit with a linear regression yielding the equation below with a strong statistical relationship (R^2^ = 0.9997):

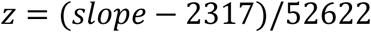

It is good to note that the charge calibration curve was specific to each instrument. Small differences in ion trajectories into the Orbitrap and differing imperfections of the Orbitrap analyzer for each instrument affect the slope of the calibration function. However, once the calibration was produced, extensive testing revealed a system did not have to be recalibrated as long as the injection energy of the ions from the C-Trap into the Orbitrap analyzer did not change.

To determine the quantized *z* with a >96% rate of correct assignment, the ion STORI slope was scored based on proximity to the nearest median charge calibration value and assigned. However, the deviation of slope values was large, such that STORI slope values were incorrectly assigned a neighboring charge state. **Supplemental Fig. 8** shows the charge assignment of approximately 800 single ion signals for the isolated myoglobin +20 charge state collected over a 2 second transient. Although all the ion signals collected were known to correspond to a +20 charge, slight variations in their STORI slopes resulted in the incorrect charge assignment of 47.5% of the ion signals collected, with the incorrect charge assignments having a Gaussian-like distribution to higher and lower charge states. A slope averaging mechanism, more commonly known as the central limit theorem, was implemented to correct the misassignments.

When STORI slope values from a known charge state were averaged, there was a reduction in charge misassignment proportional to 1/√*n*, where *n* is equal to the number of ion STORI slope values being averaged. **Supplemental Fig. 8** illustrates the implementation of this averaging mechanism, where the +20 charge assignment for the myoglobin species improved to 81.1% and 96% when averaging 4 and 16 ion STORI slope values, respectively, and subsequently assigning charge. This averaging of 16 STORI slopes was cemented into the algorithm for data acquisition utilized in the creation of all denatured hCDMS spectra in this report to accurately assign charge to individual analyte ions from 6 – 100 kDa carrying +6 to +80 charge states in the 500 – 5,000 *m/z* range. Averaged STORI slope counts increased up to 100 to assign >96% of individual ions to the correct charge state for native species in the 10,000 – 23,000 *m/z* range. Before averaging STORI slope values, first all ions collected were ordered as a function of their *m/z* values. A slope tolerance for STORI slope values 30% above or below the median slope value of the points being averaged in addition to a maximum 5 *m/z* window for ordered STORI slopes to be averaged provided strict guidelines to remove outlier values. STORI slope values are averaged pre-charge calibration and the resulting calculated *z* was parsed back to all input single ions. Once an accurate quantized charge was calculated for each single ion the mass was determined: 

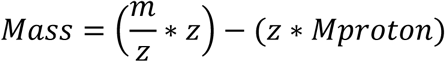

### Plotting of hCDMS Spectra (Fig. 1, Step 5)

Once the mass of each ion was calculated, the thousands of acquired individual ions were binned to create the hCDMS spectrum. Histogram bin sizes varied based on the species analyzed. Masses under 200 kDa ions were binned in 0.2 Da increments. As masses increased beyond 200 and 1,000 kDa and resolution decreased, bin sizes were increased to 30 and 500 Da, respectively. In tandem, the calculated charge of each ion utilized to create the hCDMS spectrum was binned in quantized domains to validate charge assignment. A simple protein example demonstrating these outputs is shown in **Supplemental Fig. 9. Supplemental Fig. 9a,b,c** represent the traditional *m/z* mass spectrum of myoglobin, corresponding to the hCDMS spectrum in the mass domain, and single ion charge assignment histogram for ions utilized to create the hCDMS spectrum, respectively. **Supplemental Fig. 9a** mirrors the charge state distribution in **Supplemental Fig. 9c**.

## Acknowledgements

This work was funded by the Intensifying Innovation program from Thermo Fisher Scientific and was carried out in collaboration with the National Resource for Translational and Developmental Proteomics under Grant P41 GM108569 from the National Institute of General Medical Sciences, National Institutes of Health with additional support from the Sherman Fairchild Foundation. In addition, we would like to thank Luca Fornelli and Timothy K. Toby for collecting and analyzing the HEK-293 LC/MS runs utilized for our intact mass tag hCDMS analysis.

---

**Supplemental Figure 1.**
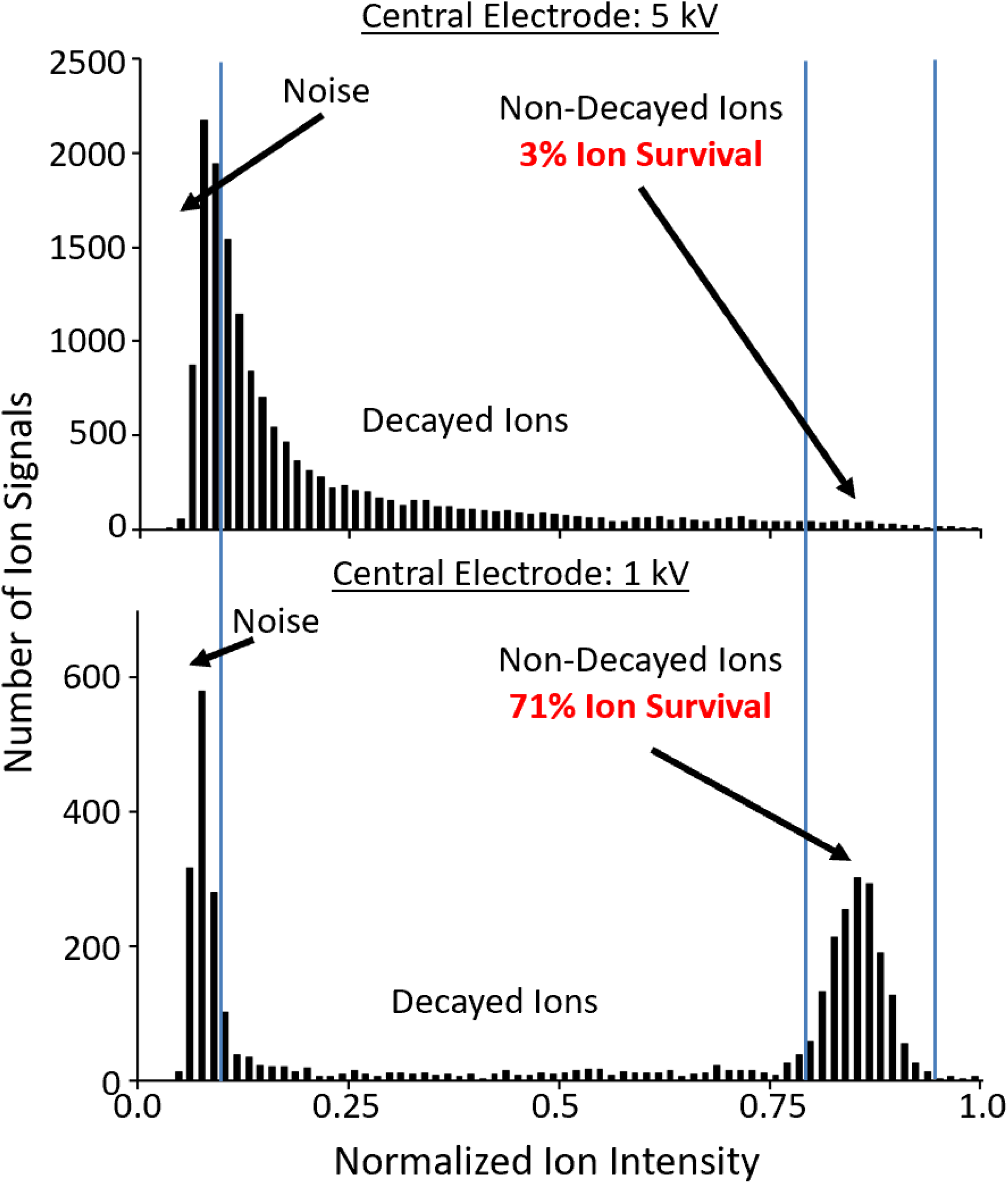
Histogram of individual ions collected that demonstrate the increased ion survival with decreased Orbitrap central electrode (CE) voltages. The solid blue lines indicate separation of low-intensity noise peaks, mid-level intensity real ions that have decayed during the acquisition event, and high-intensity real ions that did not decay during the detection event. The high intensity ions are used to create an hCDMS spectrum. As the CE voltage was decreased from −1 kV to −5 kV, ion velocity decreased and this increased ion survival over a 4 second transient from just 3% (CE = −5 kV) to 71% (CE = −1 kV). This finding drastically increased the efficiency of hCDMS during its development.

**Supplemental Figure 2.**
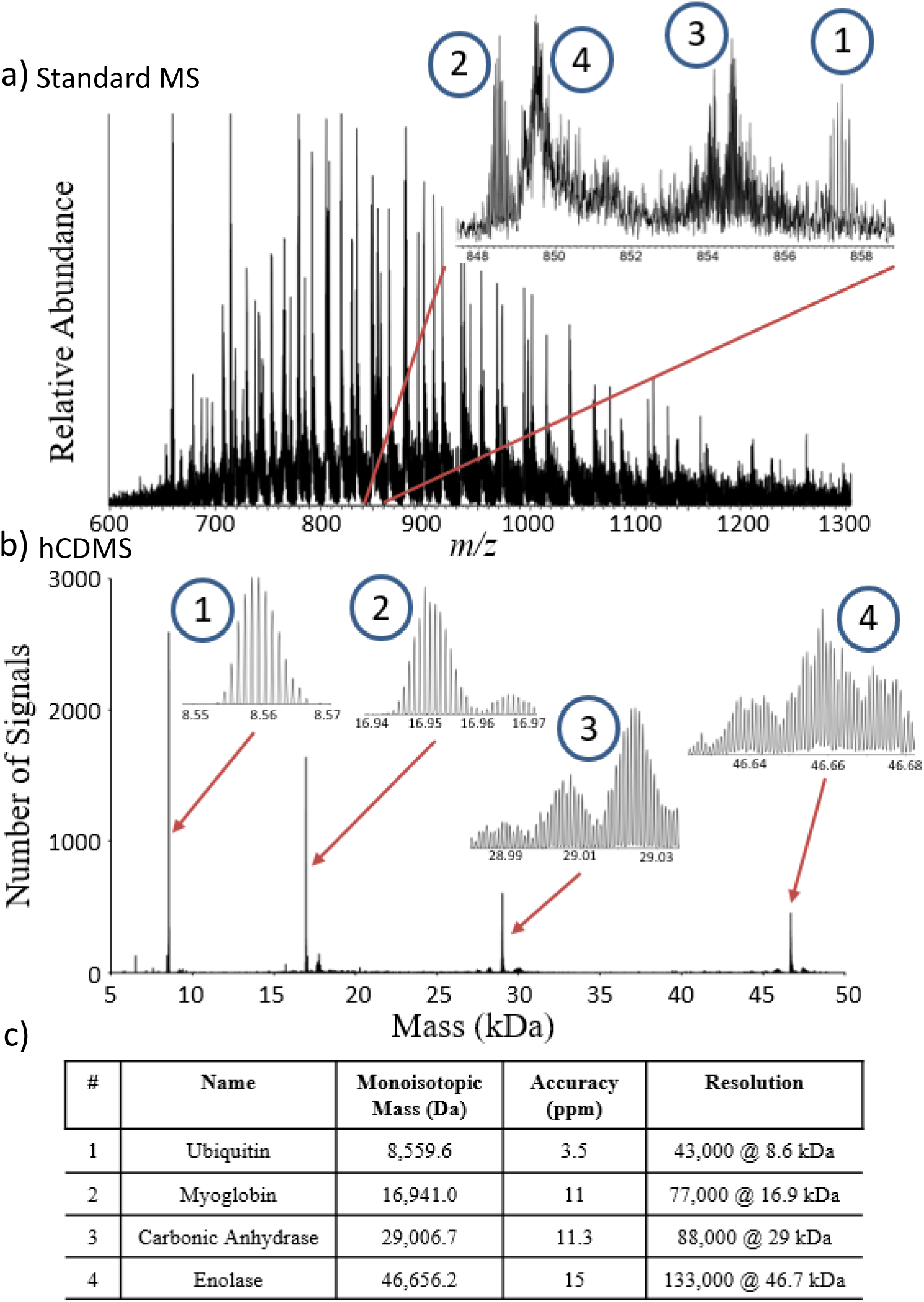
Spectra of a synthetic mixture of intact proteins created by tradition ensemble ion collection plotted in *m/z* space (a), or via the hCDMS process (b). The table in panel (c) lists the known proteins in the mixture along with their masses and corresponding figures of merit determined from the hCDMS spectrum in panel (b). The number assigned to each protein in the table correspond to the labeled features in the *m/z* spectrum and corresponding hCDMS spectrum in panels (a) and (b), respectively.

**Supplemental Figure 3.**
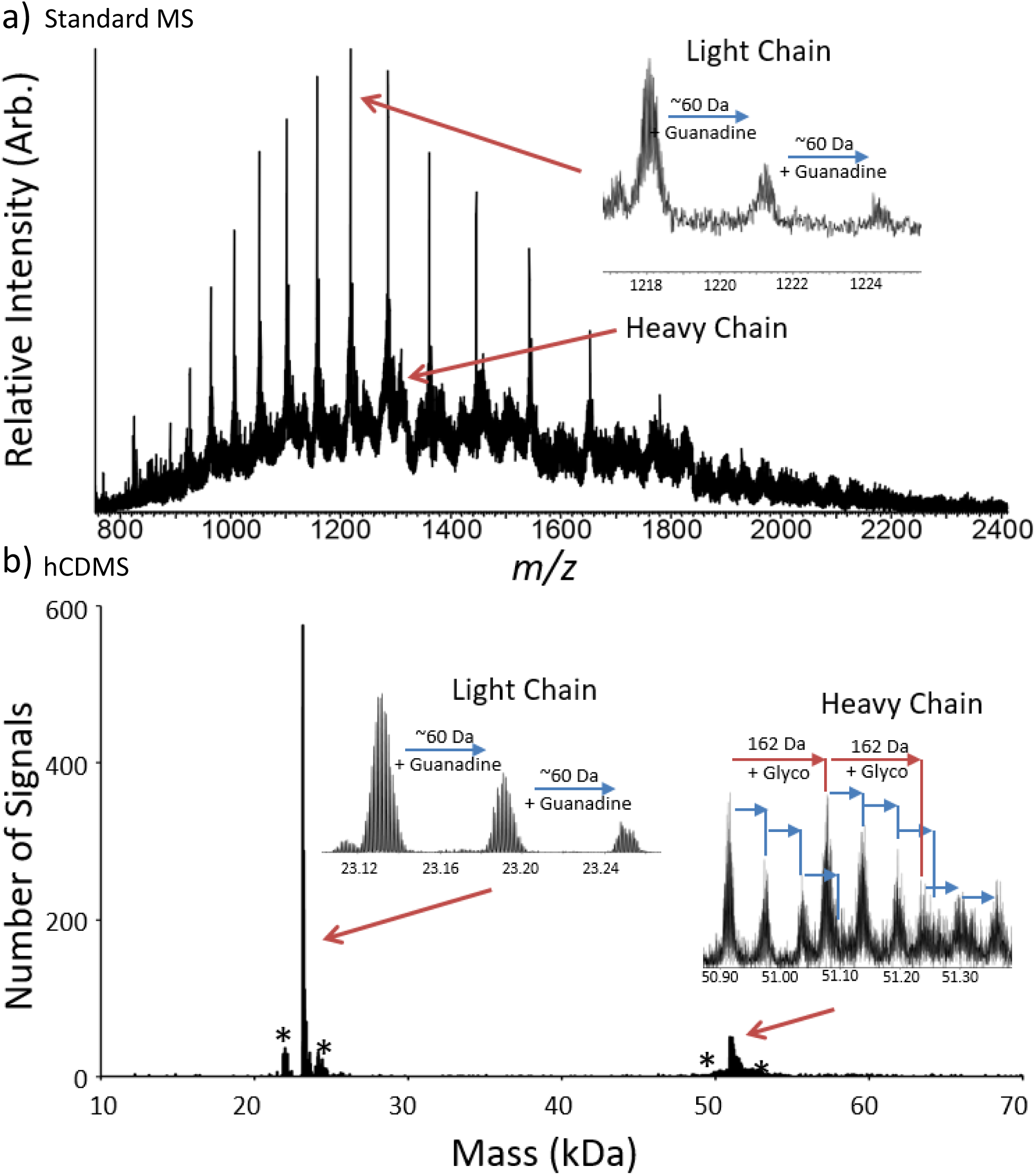
Heavy and Light chain of a monoclonal IgG measured by conventional ensemble FT-MS analysis (a) compared to hCDMS (b). Various forms of the Heavy and Light chain species including guanidine adducts and glycosylations caused a rise in baseline in the ensemble spectra were easily distinguished by hCDMS and caused no such rise in spectral baseline. Blue arrows are indicative of approximately a 60 Da guanidine adduct and red arrows indicate the addition of a 162 Da hexose-type glycosylation. Peaks arising from assignment of +1 or −1 charge state to individual ions are labeled in the hCDMS spectrum with an asterisk*.

**Supplemental Figure 4.**
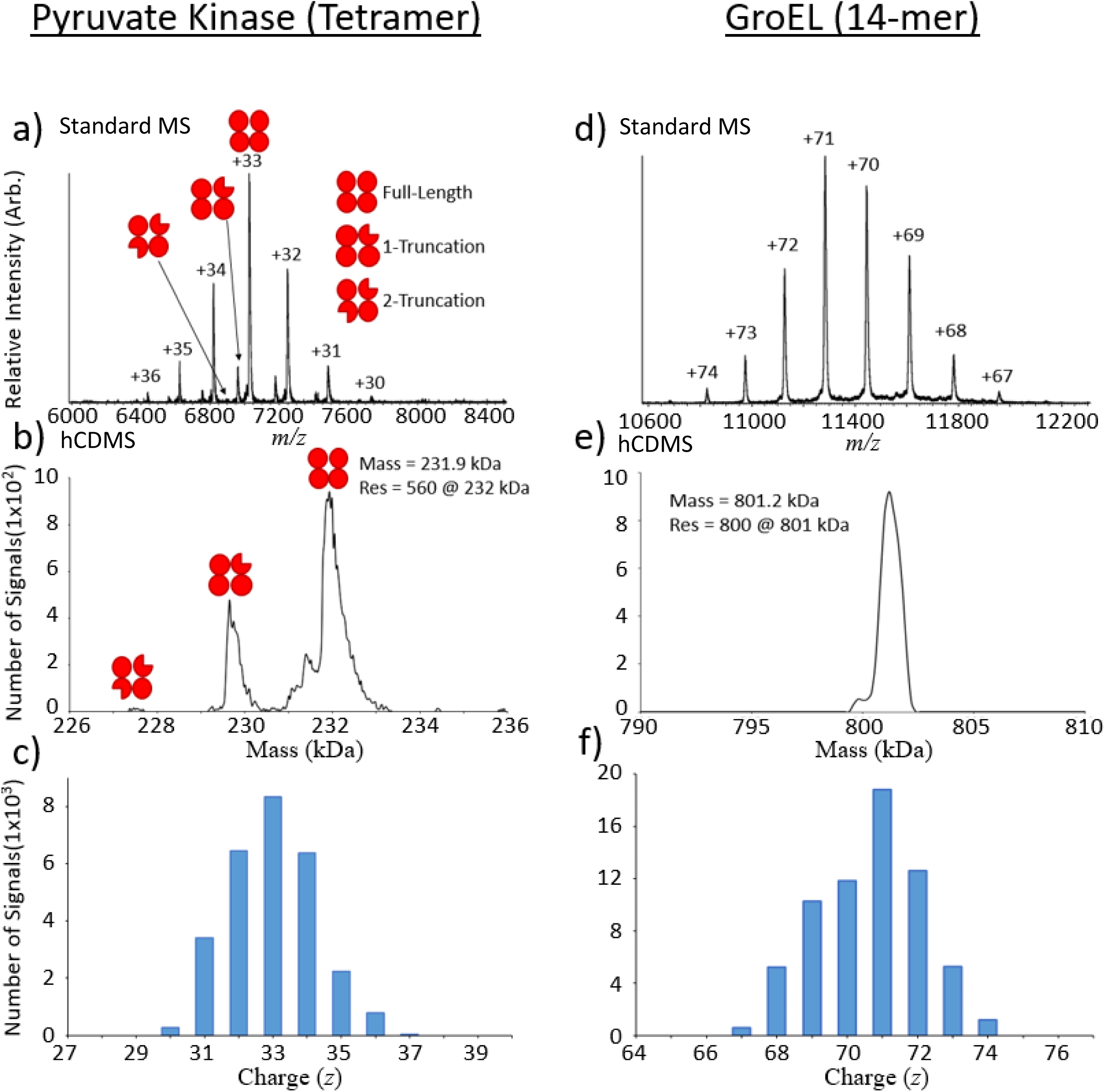
Ensemble ion spectra plotted in *m/z* space (a,d) and their corresponding hCDMS mass spectra (b,e) for the ∼232 kDa pyruvate kinase (left) and 801 kDa groEL (right) complexes with labeled charge states in panels (a) and (d). The implementation of hCDMS yields accurate spectra in the mass domain with resolution (Res) comparable to the various charge states collected in *m/z* space. Charge state values and relative charge state intensity assignments (c,f) agreed on average within <5% with their corresponding *m/z*-domain spectra (a,d).

**Supplemental Figure 5.**
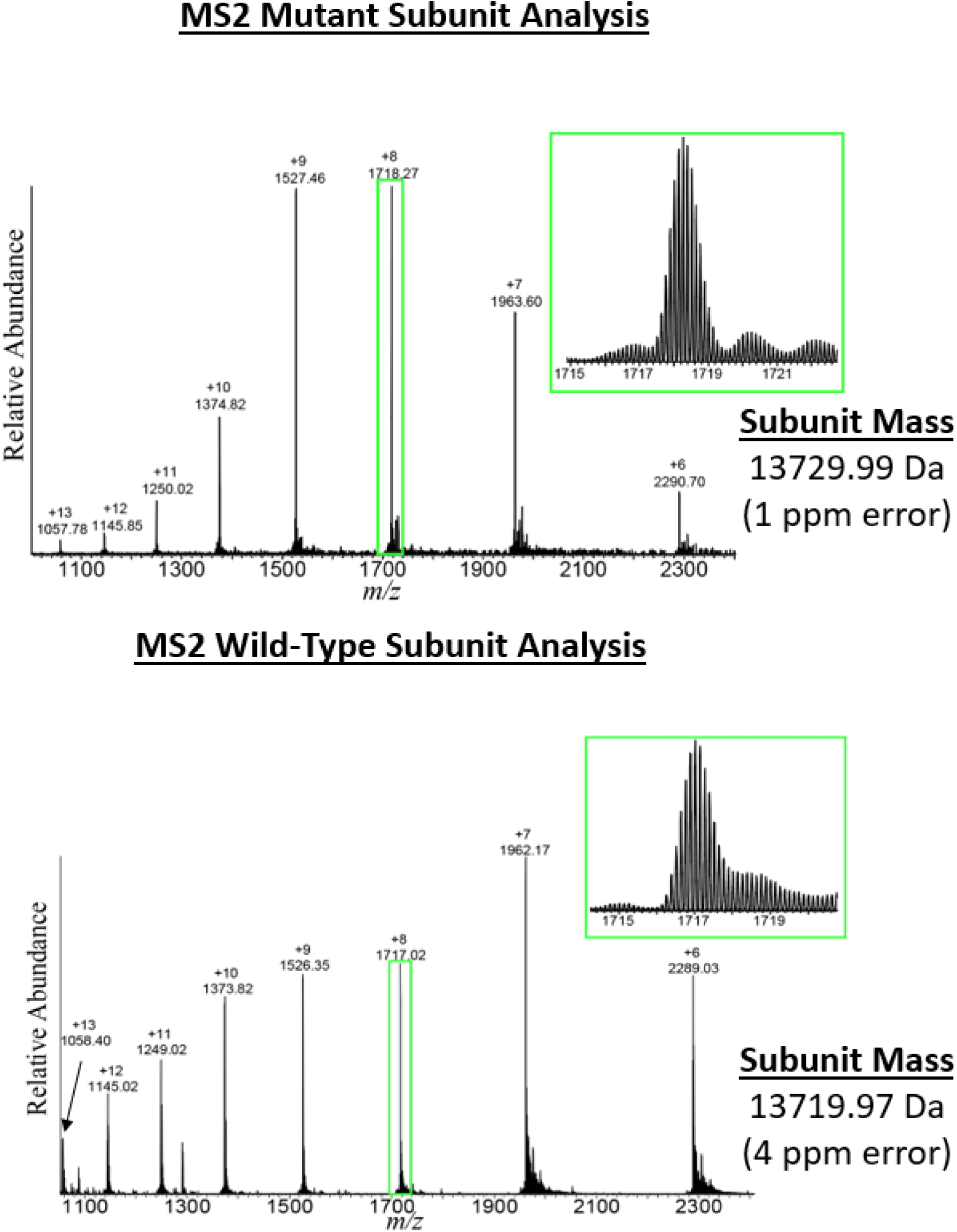
Characterization of the capsid coat proteins (CPs) used to assemble MS2 virus-like particles analyzed in **Fig. 3**. The WT (top) and MINI (bottom) CP masses are within 4 part-per-million of their theoretical values. Mass measurements from denaturing the VLP capsids produced intact mass values that match and confirm the sequence differences of each species. The one differing amino acid (proline vs. serine at position 37) between the two proteoforms results in the structural differences between the WT and MINI VLPs.

**Supplemental Figure 6.**
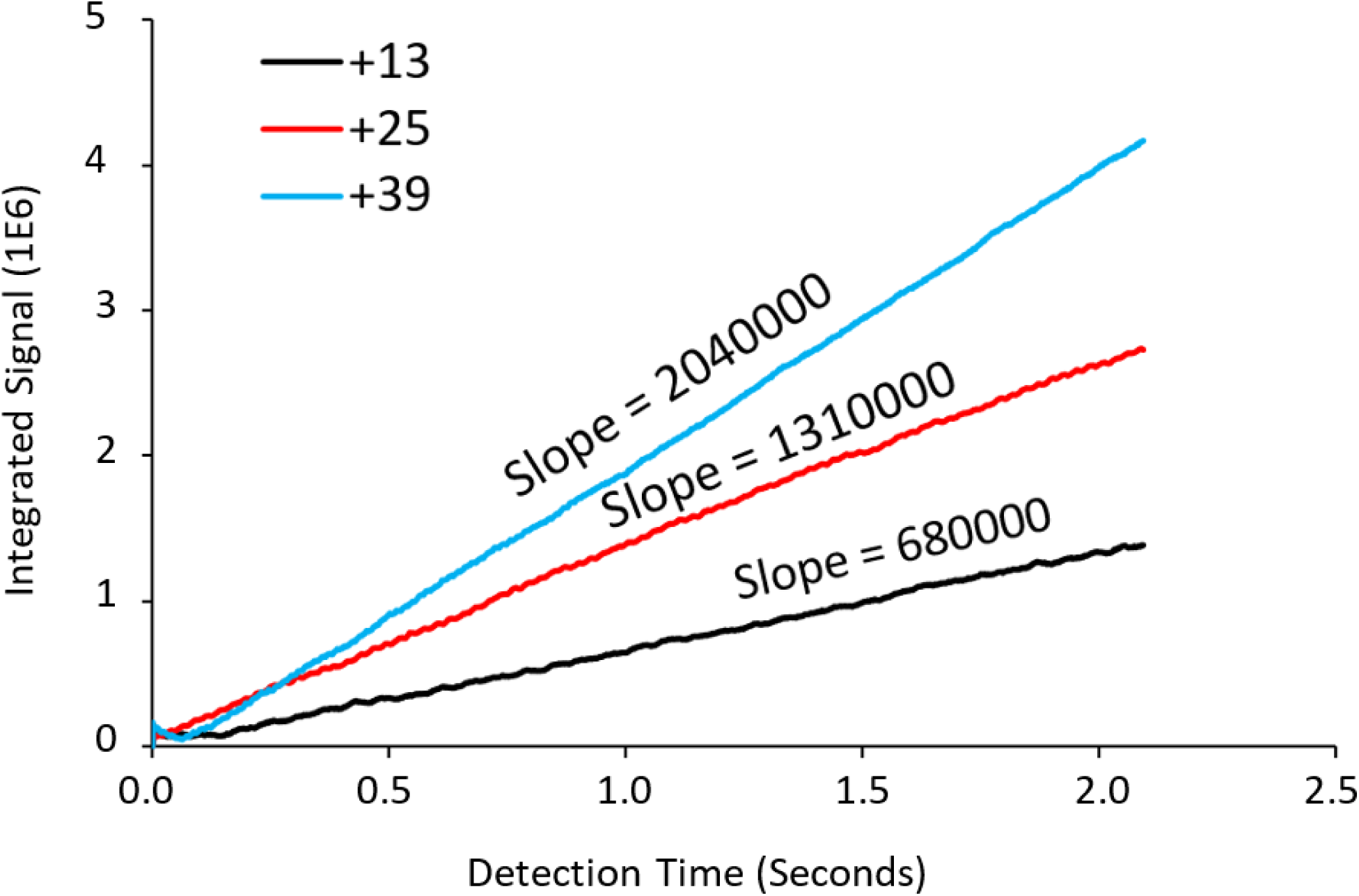
Single Ion STORI plots and their slope values for +13 (black), +25 (red), and +39 (blue) charge states of ubiquitin, myoglobin, and carbonic anhydrase, respectively. The slope of the Selective Temporal Overview of Resonant Ion (STORI) plot of an individual ion on the outer electrode of the Orbitrap is proportional to the charge of the ion signal. The STORI slope is calibrated externally to provide a single, integer result. A complete description of the STORI process will be available soon^11^ and see **Fig. 1** for an overview of the entire hCDMS process.

**Supplemental Figure 7.**
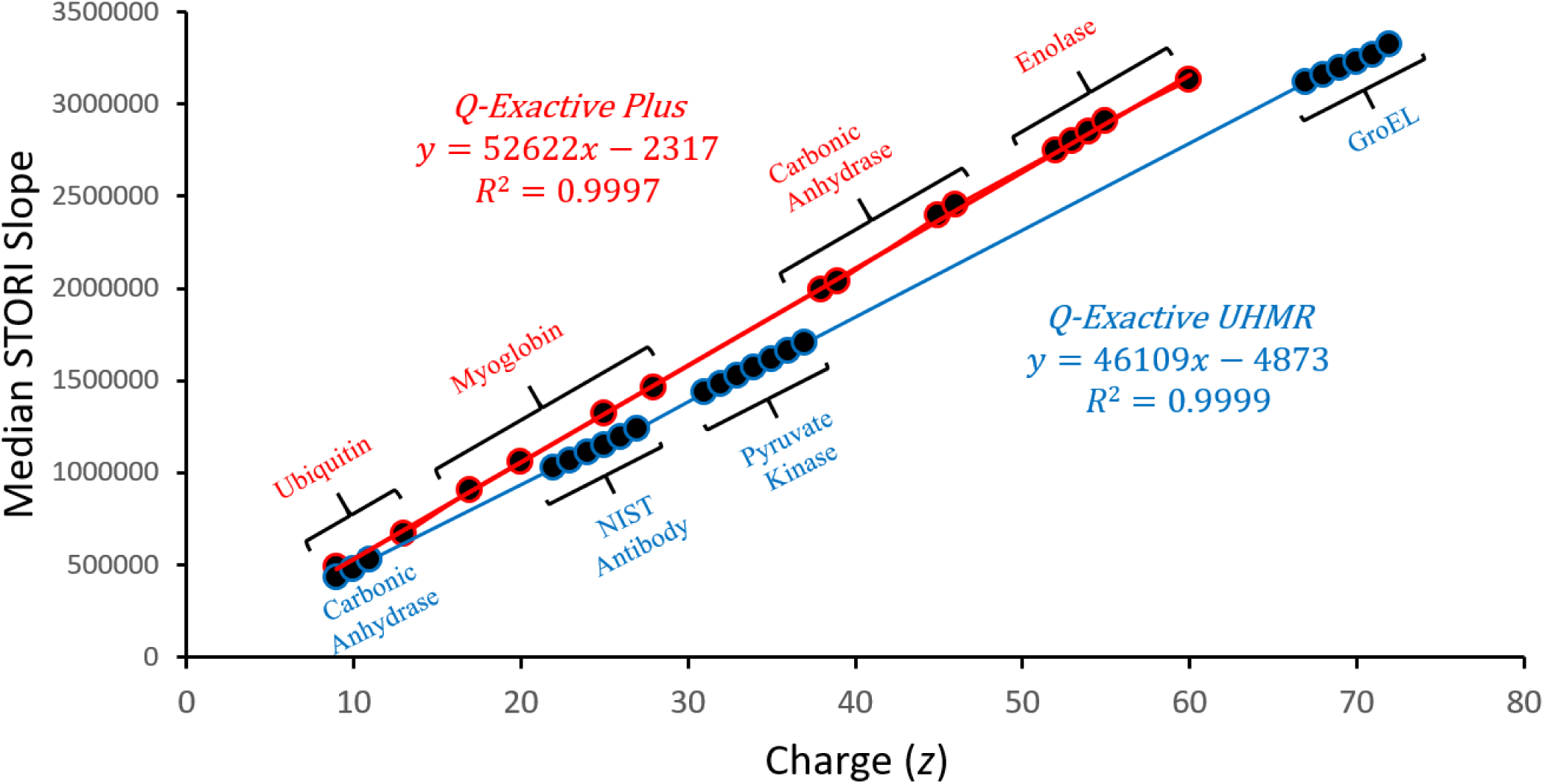
Linear calibration of STORI slope values as a function of charge state. Calibration is utilized to determine the charge of an unknown single ion signal with a calculated STORI slope. The rigorously linear statistical relationship between single ion STORI slopes and quantized charge states independent of ion *m/z* is the foundation hCDMS is built upon. The proteins utilized to produce the various charge states are labeled by their respective points on the graph. The highlighted red points/trace correspond to the calibration function on a Q-Exactive Plus instrument while the blue points/trace correspond to the calibration function on a Q-Exactive UHMR instrument.

**Supplemental Figure 8.**
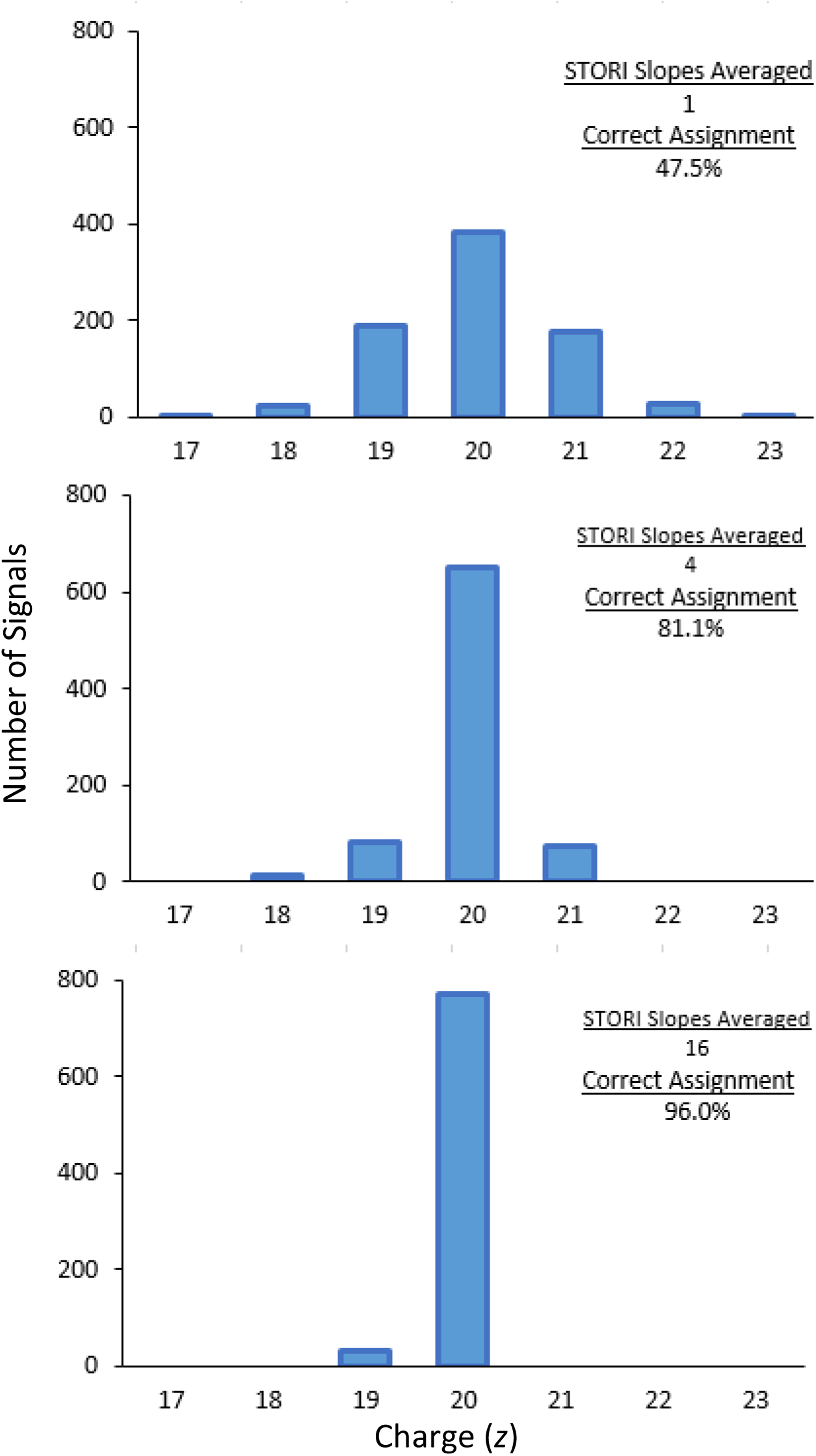
Correct assignment of known +20 myoglobin single ions. As a larger number STORI slope values are averaged prior to the assignment of an ion’s charge (top to bottom) the percentage of ions assigned to the correct (+20) charge state increases markedly. Other examples across the range of charge states used in this study give results from 94.0% to 98.5% in the rate of correct charge states for knowns analyzed by hCDMS in either denatured or native modes.

**Supplemental Figure 9.**
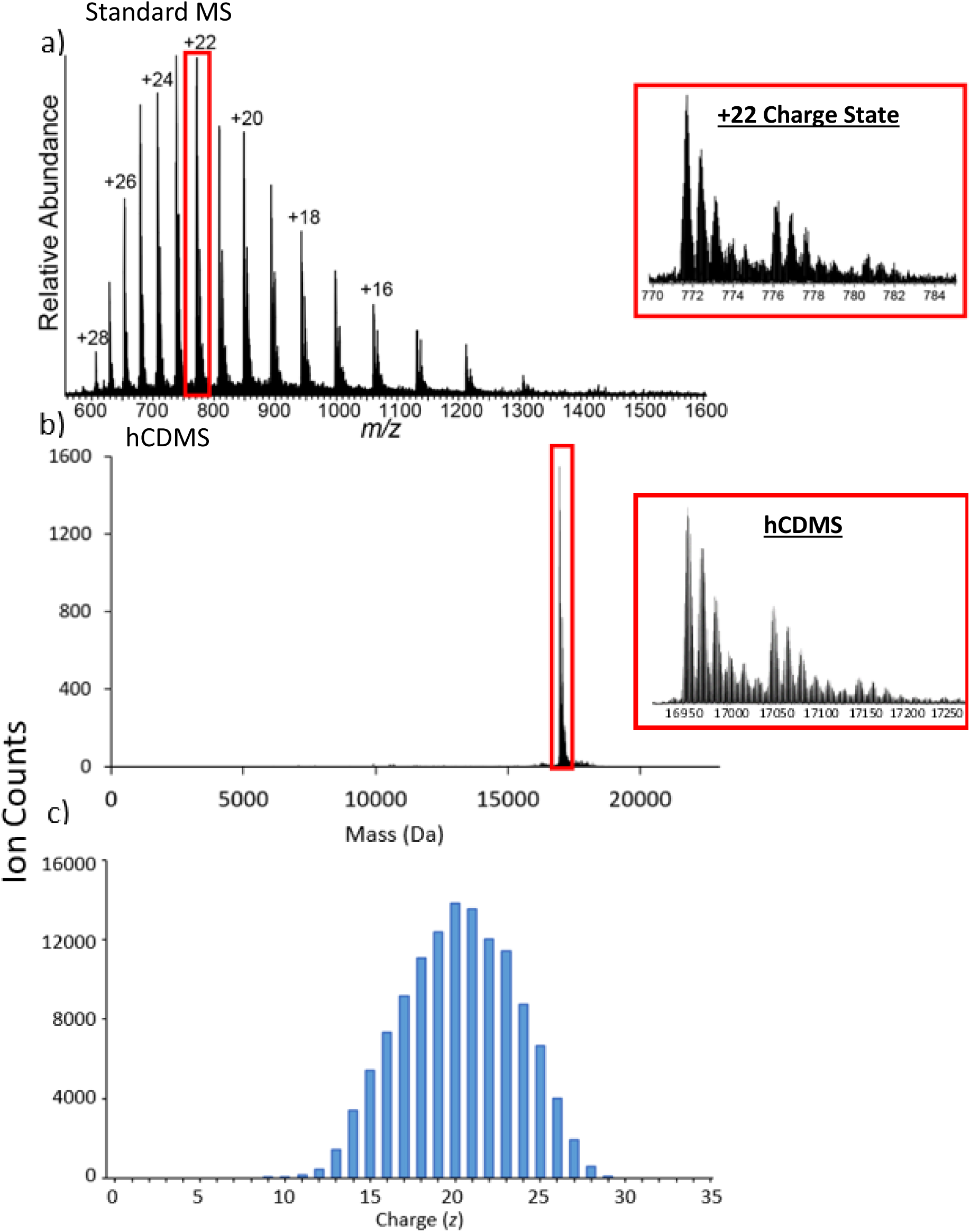
The ensemble spectrum in the *m/z* domain (a), hCDMS mass spectrum (b), and corresponding single ion charge assignment histogram (c) for the myoglobin charge state envelope. The insets in the *m/z* spectrum and hCDMS mass spectrum demonstrate that although the charges have been condensed in the hCDMS spectrum, the features within each charge state remain unchanged.

